# Deciphering cellular transcriptional alterations in Alzheimer’s disease brains

**DOI:** 10.1101/2020.04.15.041376

**Authors:** Xue Wang, Mariet Allen, Shaoyu Li, Zachary S. Quicksall, Tulsi A. Patel, Troy P. Carnwath, Joseph S. Reddy, Minerva M. Carrasquillo, Sarah J. Lincoln, Thuy T. Nguyen, Kimberly G. Malphrus, Dennis W. Dickson, Julia E. Crook, Yan W. Asmann, Nilüfer Ertekin-Taner

## Abstract

Large-scale brain bulk-RNAseq studies identified molecular pathways implicated in Alzheimer’s disease (AD), however these findings can be confounded by cellular composition changes in bulk-tissue. To identify cell intrinsic gene expression alterations of individual cell types, we designed a bioinformatics pipeline and analyzed three AD and control bulk-RNAseq datasets of temporal and dorsolateral prefrontal cortex from 685 brain samples. We detected cell-proportion changes in AD brains that are robustly replicable across the three independently assessed cohorts. We applied three different algorithms including our in-house algorithm to identify cell intrinsic differentially expressed genes in individual cell types (CI-DEGs). We assessed the performance of all algorithms by comparison to single nucleus RNAseq data. We identified consensus CI-DEGs that are common to multiple brain regions. Despite significant overlap between consensus CI-DEGs and bulk-DEGs, many CI-DEGs were absent from bulk-DEGs. Consensus CI-DEGs and their enriched GO terms include genes and pathways previously implicated in AD or neurodegeneration, as well as novel ones. We demonstrated that the detection of CI-DEGs through computational deconvolution methods is promising and highlight remaining challenges. These findings provide novel insights into cell-intrinsic transcriptional changes of individual cell types in AD and may refine discovery and modeling of molecular targets that drive this complex disease.

## Background

Alzheimer’s disease (AD) is a neurodegenerative disease that affects ∼5.7 million patients with annual cost of more than $230 billion in the US^1^. Effective disease-modifying drugs are still elusive despite the urgent need and increasing global burden^2,3^. Pathologically, AD is marked by amyloid-beta plaques and neurofibrillary tangles, along with neuronal loss and gliosis in the affected brain regions. Transcriptome-wide expression levels have been analyzed from bulk brain tissue of hundreds of AD patients and neuropathologically normal controls^4-8^ to discover genes and biological pathways that are perturbed in and/or lead to AD. Systems biology and bioinformatics analysis of these data have implicated altered pathways in AD including immune response ^6^ and myelin metabolism ^4,5^. However, a fundamental knowledge gap remains concerning whether disease-associated changes in brain gene expression are due to changes in cellular composition of the AD samples secondary to disease neuropathology, or due to changes in the intrinsic regulation/activity of genes in the central nervous system (CNS) cells. From a clinical perspective, it is difficult to target changes in cellular composition secondary to neuropathology, whereas intrinsic changes in gene expression that may drive AD progression are potentially “druggable”.

We expect that identifying cell-intrinsic differentially expressed genes (CI-DEGs) of individual CNS cell types will reveal novel insights into the genes and pathways that could potentially identify drug targets and lead to AD therapeutics. This approach circumvents the influence of cell-composition changes that can impact disease associated DEGs obtained from bulk tissue transcriptome analysis. RNA sequencing (RNAseq) studies from single cell, single nucleus or purified adult human CNS cells^9-11^ can be used to identify CI-DEGs. Even though such single cell-type RNAseq data can effectively serve as a reference to annotate bulk-tissue transcriptome data^4^, such approaches remain costly, cumbersome and limited in sample sizes. On the other hand, there exist large-scale bulk brain RNAseq datasets^5,8,12^, which can be leveraged to identify CI-DEGs through analytic deconvolution of bulk tissue expression into signals of individual cell types by using cell proportions or their proxies^13,14^.

In this study, we describe the design and application of a bioinformatics pipeline that uses cell type marker genes to estimate cell proportion^15,16^ to deconvolute bulk tissue transcriptome data using three computational approaches^13,14,17^ and to subsequently identify CI-DEGs. We applied our pipeline to the analysis of three post-mortem brain datasets, one from dorsolateral prefrontal cortex (DLPFC)^8^ and two from temporal cortex (TCX)^4,12,18^ regions, comprised of 685 unique samples. Consensus CI-DEGs common to both TCX and DLPFC regions were analyzed for enrichment of gene ontology (GO) terms. We compared the results of consensus CI-DEGs to consensus bulk-DEGs. In addition, for the DLPFC^8^ dataset that had both bulk and single nucleus RNAseq^19^ (snRNAseq) data, we compared the CI-DEGs from the computational deconvolution to CI-DEGs obtained from snRNAseq^19^.

To our knowledge, this is the first study to detect consensus CI-DEGs and their enriched gene ontology (GO) terms from multiple brain regions using multiple computational deconvolution algorithms for AD and control RNAseq samples. Our study illustrates the cell proportion landscape of AD and control brain regions assessed in three independent RNAseq studies^4,7,8,12^. We identify consensus CI-DEGs many of which are not observed in bulk-DEG analysis and characterize their cell-type specificity. GO terms that are enriched for CI-DEGs implicate cell intrinsic transcriptional alterations that may influence AD, rather than be a result of cell-proportion changes in this disease. These CI-DEGs and their biological pathways may serve as refined molecular targets for therapeutic discoveries and disease modeling in AD. Our study also demonstrates that detection of CI-DEGs through computational deconvolution methods is promising while some challenges remain.

## Results

### Cellular composition in three brain cohorts from two brain regions

We analyzed three cohorts each consisting of post-mortem brains from AD and control subjects (**Table S1**), namely the Rush Religious Orders Study and Memory and Aging Project dorsolateral prefrontal cortex (DLPFC)^7,8^, Mayo Clinic temporal cortex (TCX-Mayo)^4,12^, and Mount Sinai VA Medical Center Brain Bank temporal cortex (TCX-MSBB)^18^. We generated the TCX-Mayo RNAseq dataset, and downloaded DLPFC and TCX-MSBB RNAseq datasets from the AMP-AD knowledge portal on Synapse (www.synapse.org).

Cell proportions (**Table S2**) were estimated for DLPFC, TCX-Mayo and TCX-MSBB datasets independently using the digital sorting algorithm (DSA) method^16^ and the top 100 marker genes (**Table S3**) obtained from R package BRETIGEA^15^ for each of the following cell types – neuron, oligodendrocyte, microglia, oligodendrocyte progenitor cell (OPC), astrocyte and endothelial cell.

An inspection about the pairwise correlation between marker genes (**Fig 1a**) revealed that markers of OPC have poor median pairwise Pearson correlation values of 0.12 in DLPFC, 0.11 in TCX-Mayo and 0.06 in TCX-MSBB respectively, whereas among the other five cell types neuronal markers have the highest median correlation (0.68 in DLPFC, 0.78 in TCX-Mayo and 0.67 in TCX-MSBB), and microglia markers have the lowest correlation (0.37 in DLPFC, 0.42 in TCX-Mayo and 0.44 in TCX-MSBB). In addition, a computer simulation study (**Fig S1**) demonstrated that the estimated proportions of OPC were not robust upon using different selection of marker genes. Therefore, we did not include OPC in downstream analyses in this study.

**Fig.1:**
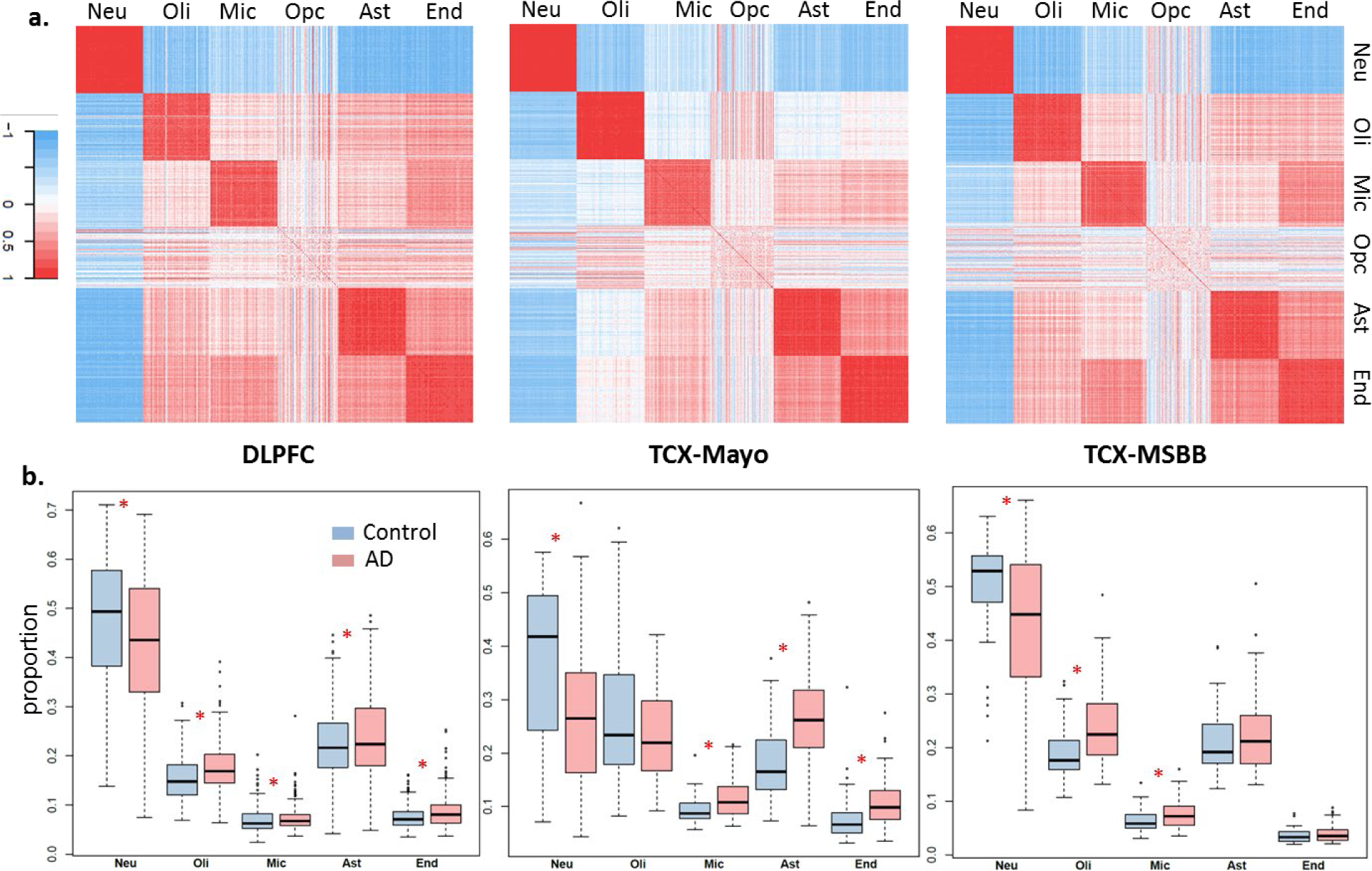
**a)** Pearson correlation between marker gene expressions in six cell types. Marker genes are from literature. **b)** Estimated cell proportions in DLFPC, TCX-Mayo and TCX-MSBB datasets in five cell types. Red asterisk indicates differences between cell proportions in AD and control groups at nominal p value 0.05 from Wilcoxon rank sum test.

In all three datasets, neuronal cell proportion estimates were significantly lower in AD compared to controls (**Fig 1b**). The magnitude of this decrease was the greatest for TCX-Mayo (AD mean proportion = 28.0%, Control = 35.7%; ratio of AD:control cell proportions=0.78), followed by TCX-MSSM (AD = 42.3%, control = 49.3%; ratio=0.87) and DLPFC (AD = 42.4%, control = 47.4%; ratio=0.89). The estimated proportions of microglia were significantly higher in AD vs. controls for all datasets, with higher magnitude in TCX-Mayo (AD:control ratio=1.19) and TCX-MSBB (AD:control ratio=1.19) than for DLPFC (AD:control ratio=1.06). The estimated proportions of astrocytes and endothelial cells were significantly higher in AD vs. controls for DLPFC and TCX-Mayo datasets, although the magnitude was greater in TCX-Mayo (1.40 and 1.30 respectively) than in DLPFC (1.07 and 1.14 respectively) for both cell types. Oligodendrocyte proportion is significantly higher in AD in DLPFC with AD:control ratio 1.14 and TCX-MSBB with AD:control ratio 1.27, although remains unchanged in TCX-Mayo with the ratio 0.94. Collectively, these findings demonstrate that the proportions of CNS cell types are different in post-mortem AD vs. control brains for most cell types. Although these proportional changes with AD are mostly consistent across the different studies, their extent varies across brain regions, with TCX tending towards higher magnitude of neuronal loss and microglia proliferation than DLPFC. It needs to be emphasized that the cell proportion changes estimated here are relative values, rather than absolute cell proportion changes between ADs and controls.

### Differential Expression Analyses

In this study, three computational approaches were applied to identify cell intrinsic differential expression in individual cell types (CI-DEGs, **Table S4-S6**), namely CellCODE^14^, PSEA^13^ and our method WLC. Differentially expressed genes from bulk brain tissue (bulk-DEGs) were identified through linear regression without adjusting for cellular composition (**Table S7**). For the DLPFC, TCX-Mayo and TCX-MSBB datasets, we obtained bulk-DEGs and CI-DEGs from the three computer algorithms for neuronal, oligodendrocytic, microglial, astrocytic and endothelial cell types respectively.

We compared bulk-DEGs across the three datasets (**Fig 2a**, top panel). Similarly, CI-DEGs from CellCODE, PSEA and WLC are compared across datasets (**Fig 2a**, lower panels), such that CI-DEGs shared between datasets are required to be consistent in the designated cell type. All DEGs are identified at nominal p-value cutoff 0.05 and shared CI-DEGs have the same direction of change in the compared datasets. The ratio of overlap between any two datasets over all DEGs, i.e. the number in overlapping areas of the Venn diagram over the total number (**Fig 2a**, top panel), is 30.0% or 1711/5697 in up-regulated bulk-DEGs, and 34.8% or 2214/6371 in down-regulated bulk-DEGs. This ratio of overlap in bulk-DEGs is much higher than that in CI-DEGs (2.7%, 4.7% and 10.0% in up-regulated genes from CellCODE, PSEA and WLC respectively; 3.1%, 6.8% and 9.3% in down-regulated genes from CellCODE, PSEA and WLC respectively).

**Fig.2:**
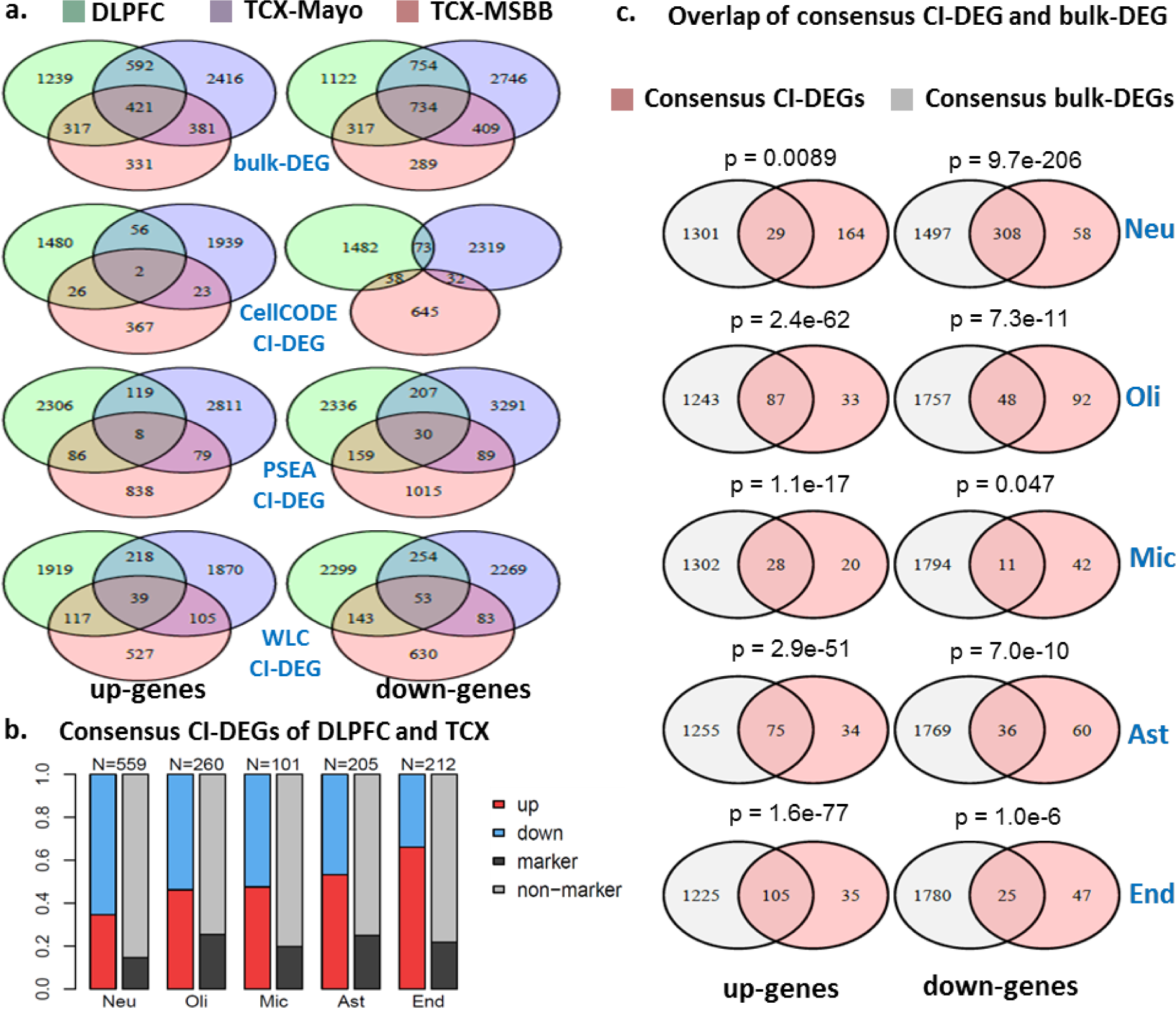
**a)** Overlap across three independent RNAseq datasets of bulk-DEGs (upper panel) and CI-DEGs (lower panels) from three computational approaches. **b)** Consensus CI-DEGs between DLPFC and TCX brain regions, which consist of consensus CI-DEGs between DLPFC and TCX-Mayo, or between DLPFC and TCX-MSBB. c) Overlap between consensus CI-DEGs and consensus bulk-DEGs, per cell type. The p-values of overlap are from hypergeometric tests.

### Consensus CI-DEGs between DLPFC and TCX

To obtain the consensus CI-DEGs that are shared between DLPFC and TCX brain regions, we selected those CI-DEGs that are detected in “DLPFC and TCX-Mayo” or in “DLPFC and TCX-MSBB” under any of the three algorithms (**Fig S2**). We combined all such genes, which collectively comprised the consensus CI-DEGs for each cell type (**Fig 2b**). Similarly, consensus bulk-DEGs were the combined set of bulk-DEGs shared between “DLPFC and TCX-Mayo” or “DLPFC and TCX-MSBB”.

Most consensus CI-DEGs are from neuronal cells (N=559), followed by oligodendrocytes (N=260), whereas microglia contributed the least number (N=101). The majority (65.5% or 366/559) of neuronal CI-DEGs is down-regulated in AD, and the majority (66.0% or 140/212) of endothelial CI-DEGs is up-regulated in AD, with other cell types lying in between. Some of these CI-DEGs are also among the 1000 marker genes of the corresponding cell type from BRETIGEA^15^; 14.7% or 82/559 of neuronal CI-DEGs are also neuronal markers, 25.4% or 66/260 of oligodendrocyte CI-DEGs are also oligodendrocyte markers, and other cell types lie in between.

With regards to consensus bulk-DEGs (**Fig S3**), 28.2% of them (885/3135) are cell type markers; 10.4% neuronal markers, 5.6% oligodendrocyte, 3.4% microglia, 4.8% astrocyte and 4.0% endothelial markers. The above observations indicate that computational deconvolution algorithms could identify CI-DEGs for both marker genes and non-marker genes. Importantly, the proportion of non-marker CI-DEGs is greater than that in bulk-DEGs. This suggests that compared to bulk-DEGs, CI-DEGs may be capturing a greater proportion of expression changes that are not due to mere cell population changes.

We also compared the consensus bulk-DEGs with consensus CI-DEGs (**Fig S4**). We determined that only a small fraction (15.0% or 29/193) of the up-regulated neuronal CI-DEGs was also present in up bulk-DEGs although the overlap is still significant (**Fig 2c**). In comparison, most of the up-regulated CI-DEGs of the other four cell types were included in up bulk-DEGs. On the other hand, most (84.2% or 308/366) of the down-regulated neuronal CI-DEGs were also present in down bulk-DEGs, whereas most of the down-regulated CI-DEGs of the other four cell types were absent from this group. Since bulk-DEGs did not adjust for neuronal loss and gliosis in AD (**Fig 1b**), its ability to identify up-regulated neuronal genes and down-regulated glial genes is likely to be compromised. For the same reason, bulk-DEGs may have a false inflation of detecting down-regulated neuronal and up-regulated glial genes.

### Enriched GO terms of consensus CI-DEGs between DLPFC and TCX

To identify pathways implicated by CI-DEGs that are robust across brain regions, we performed Gene Ontology (GO) enrichment analysis^20,21^ for the consensus CI-DEGs, assessing separately those that are up vs. down in AD subjects (**Table S8-S17**). **Fig 3** illustrates the top two enriched GO terms by enrichment p-values, after filtering out terms that encompass less than four CI-DEGs or are cellular compartments.

**Fig 3:**
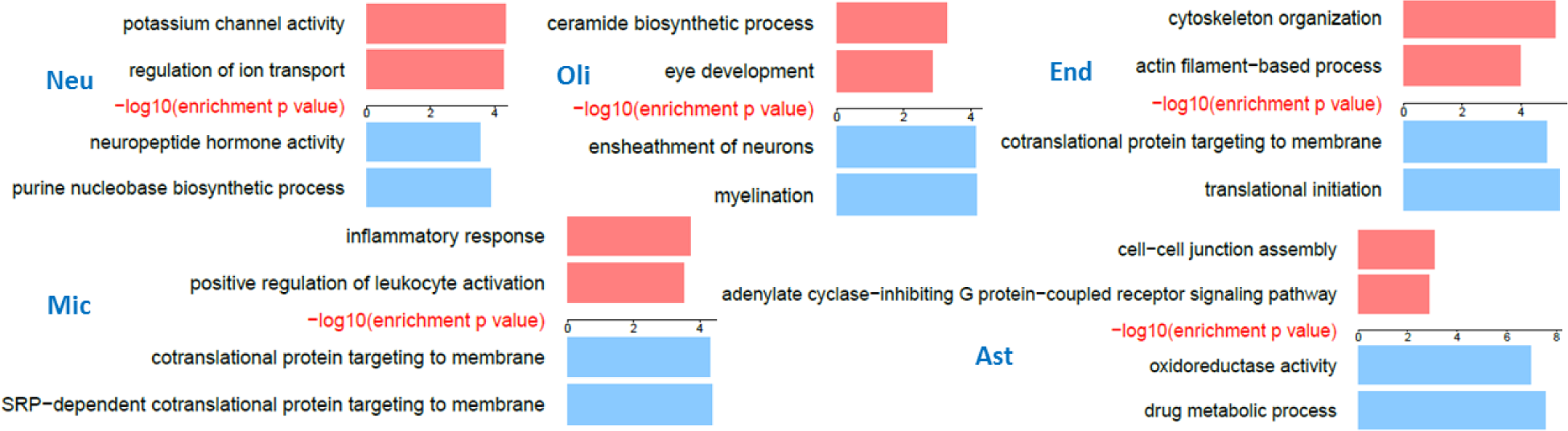
Top two enriched GO terms in up (red) or down-regulated (blue) consensus CI-DEGs between DLPFC and TCX regions, per cell type.

Consensus CI-DEGs revealed biological pathways that are perturbed in AD in specific brain cell types. Some of these pathways have previously been implicated in AD and others are novel. Down-regulated neuronal CI-DEGs were enriched in neuropeptide hormone activity (GO:0005184) and hormone activity (GO:0005179) pathways, which include *VGF* (a.k.a. neuroendocrine regulatory peptide 1)^22^ and corticotropin releasing hormone (*CRH*)^23^ (**Table S13**). Consensus up-regulated neuronal CI-DEGs were significantly enriched in potassium channel activity (GO:0005267) and regulation of ion transport (GO:0043269) pathways (**Table S8**). The latter GO term encompasses most of the genes from the former, and also includes other genes involved in neuronal functions such as the glutamate ionotropic receptor NMDA type subunit 1, *GRIN1*^*24*^ and *SYT1*, which encodes the synaptic vesicle protein, synaptotagmin^25^.

Many of the most significant GO terms are related to key functions of the respective cell types for the glial CI-DEGs, as well. The top enriched pathway of down-regulated CI-DEGs in oligodendrocytes is myelination (GO:0042552), including myelin basic protein (*MBP*)^4^, plasmolipin (*PLLP*)^4,5^, myelin and lymphocyte protein (*MAL*), and myelin-associated glycoprotein (*MAG*)^26^ (**Table S14**). Up-regulated CI-DEGs of oligodendrocytes are enriched in ceramide biosynthetic process (GO:0046513) including ceramide synthase 4 (*CERS4*) and UDP glycosyltransferase 8 (*UGT8*)^5^ (**Table S9**). Ceramide is a constituent of sphingomyelin, a sphingolipid which is particularly found in the myelin sheath; and also a multi-functional signaling molecule^27,28^. Hence, both the down-regulated and the up-regulated oligodendroglial consensus CI-DEGs highlight different components of the myelin biology that are perturbed in AD.

Similarly, microglial, astrocytic and endothelial CI-DEGs also highlight processes pertinent to the functions of these cell types. Microglial up-regulated CI-DEGs are enriched in inflammatory response (GO:000695) and leukocyte activation (GO:0002696), which includes complement C3a receptor 1 (*C3AR1*)^29^, interleukin 18 (*IL18*)^30,31^ and CCAAT enhancer binding protein alpha (*CEBPA*)^32^ genes (**Table S10**).

Astrocytes, a cell type that plays a critical role in maintaining brain energy dynamics^33^ and metabolism^34^, show enrichment of oxidoreductase activity (GO:0016491) and drug metabolic process (GO:0017144) in down-regulated CI-DEGs which includes genes glutathione S-transferase mu 2 (*GSTM2*)^35^ and thioredoxin2 (*TXN2*)^36^ (**Table S16**). Astrocytic up-regulated consensus CI-DEGs are enriched for cell-cell junction assembly (GO:0007043) process (**Table S11**), including the astrocytic gap junction protein connexin43 (*GJA1*)^37^, which was identified as a key regulator associated with AD related outcomes. The other top GO process for astrocytic up-regulated consensus CI-DEGs is adenylate cyclase-inhibiting G protein-coupled receptor signaling pathway (GO:0007193), which harbors adenylate cyclase 8 (*ADCY8*), involved in memory functions^38^.

Finally, endothelial cells, which are crucial in maintaining blood-brain barrier integrity^39,40^, show enrichment of up-regulated DEGs in cytoskeleton organization (GO:0007010) and actin filament-based process (GO:0030029) (**Table S12**).

Importantly, some CI-DEGs highlight protein translation as a top perturbed biological pathway. Down-regulated microglial consensus CI-DEGs show enrichment in processes involved in protein translation (GO:0006614 and GO:0006613), which include ribosomal protein encoding genes^41-43^ (**Table S15**). Similarly, down-regulated endothelial consensus CI-DEGs also harbor ribosomal protein encoding genes, with enrichment in protein translation related GO processes (GO:0006413 and GO:0006613) (**Table S17**).

### Comparison of CI-DEGs from computational deconvolution vs. snRNAseq

We determined the extent to which each of the three computational deconvolution algorithms could detect CI-DEGs from bulk tissue by comparison of their results with those obtained in a published snRNAseq study^19^. The ROSMAP dataset utilized in our study has both bulk RNAseq from DLPFC (bulk-DLPFC) as well as snRNAseq (snDLPFC) in a subset of its participants^19^. We compared the bulk-DLPFC data deconvoluted with three different algorithms with the published snDLPFC^19^ data. Endothelial CI-DEGs were not available from the snRNAseq study, therefore overlap of results could be assessed only for four cell types.

We tested the overlap between the top CI-DEGs for each cell type obtained from deconvoluted bulk-DLPFC and those from snDLPFC ranked by their p values (**Fig 4a**). We evaluated the overlap for a range of top CI-DEGs up to top 1,000 genes. Overlap for CI-DEGs that are either up (**Fig 4a, upper panel**) or down (**Fig 4a, lower panel**) in AD were assessed separately. Hence, overlapping genes had both similar ranks and direction of effect in both deconvoluted bulk-DLPFC and snDLPFC analyses. We established the significance of overlap using simulations for a range of top ranked genes (N=200, 600 and 1,000) (**Table S18**).

**Fig 4:**
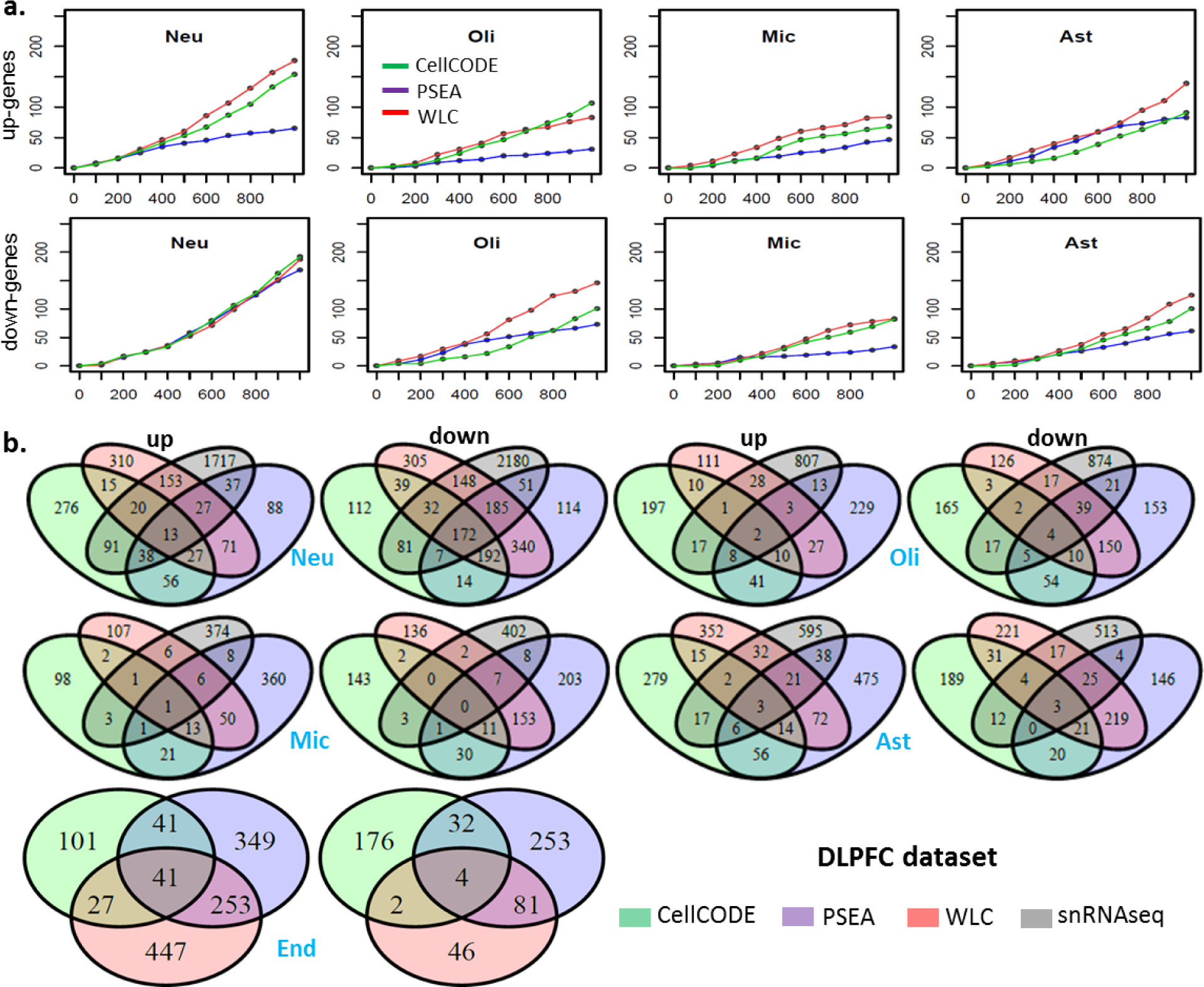
Comparison of CI-DEGs from computational deconvolution with CI-DEGs from snRNAseq on DLPFC dataset. **a.** Upper panel: number of overlapping genes (y-axis) between the top *N* (x-axis) up-regulated genes in snDLPFC and top *N* up-regulated genes from bulk-DLPFC deconvoluted with PSEA, WLC and CellCODE, respectively. Lower panel: number of overlapping genes (y-axis) between the top *N* (x-axis) down-regulated genes in snDLPFC and top *N* down-regulated genes from deconvoluted bulk-DLPFC. **b.** Venn diagram of CI-DEGs from computational deconvolution methods and those from snRNAseq. Overlap is evaluated for both bulk-DLPFC and snDLPFC CI-DEGs detected at nominal p value ≤0.05.

Neuronal CI-DEGs retained their significance of overlap across all comparisons and for all algorithms, except for the top 1,000 up-regulated neuronal CI-DEGs deconvoluted with PSEA. Microglial CI-DEGs had the least numbers of significant overlap for their top ranked genes. Astrocytic and oligodendrocytic top ranked CI-DEGs had significance of overlap between the neuronal and microglial results (**Table S18**). These findings are reflective of the abundance of these cell types, with the most abundant neurons having the most overlap for the top ranked CI-DEGs between deconvoluted bulk-DLPFC and snDLPFC.

Amongst these comparisons, we determined that the significance for overlap was best for all algorithms for the top ranked 600 genes. Using WLC deconvoluted results, the overlap for the top 600 CI-DEGs from bulk-DLPFC and snDNPFC are statistically significant for all eight comparisons (**Fig 4a**). For the top 600 genes, overlap with CellCODE results is significant for all except down-regulated oligodendrocyte and up-regulated astrocyte CI-DEGs. For PSEA, none of the microglia CI-DEGs had significant overlap. PSEA results for the top 600 genes were otherwise significant for all but up-regulated oligodendrocyte and down-regulated astrocyte genes.

We also performed a comparison of CI-DEGs identified at nominal significance (p-value <0.05) with each algorithm from bulk-DLPFC to nominally significant snDLPFC results (**Fig 4b, Table S19**). As with the above comparison, genes that are either up or down in both deconvoluted bulk-DLPFC and snDLPFC data were analyzed separately for each cell type.

Not surprisingly, down-regulated neuronal CI-DEGs have the greatest overlap (537/3732 or 14.4% for WLC, 292/3213 or 9.1% for CellCODE, 415/3516 or 11.8% for PSEA). These overlaps are significant for all three algorithms (**Table S19**). Down-regulated CI-DEGs in microglia show the least proportion of overlap (9/723 or 1.2% for WLC, 4/609 or 0.66% for CellCODE, 16/820 or 2.0% for PSEA) (empirical p-value > 0.05). Significant overlap detected with WLC (all but down-regulated microglia) and PSEA (all but microglial results and up-regulated oligodendrocytes) were similar, whereas CellCODE results had significant overlaps only for the neuronal CI-DEGs (**Table S19**).

## Discussion

There is an increasing number of large scale RNAseq-based transcriptome datasets generated in bulk tissue for many diseases, including brain tissue from patients with AD, other neurodegenerative diseases and controls^5-8,12^. Comparison of such transcriptome data from patient and control individuals has been instrumental in the identification of genes and co-expression networks that are altered in and may therefore be risk factors for these diseases^4-7,44^. The discovery that many of these transcriptional networks harbor genes with disease risk variants provides support for the utility of this bulk-transcriptome approach in deciphering molecules and pathways that are risk factors for these conditions. Nevertheless, there is clear evidence for abundant transcriptome alterations in bulk tissue from affected organs of patients with disease, which appears to be due to changes in cellular composition of that tissue as a consequence of the disease processes^4,45^. Given this, there is a strong need to detect cell-intrinsic transcriptional alterations to be able to distinguish gene expression changes that are upstream of and may therefore be causative factors for disease from those that are merely a result of cell proportion changes that occur due to disease pathology. The discovery of molecules and pathways that are upstream of and risk factors for disease pathology is of paramount importance for development of targeted therapies. This information can also aid in the identification of refined disease biomarkers reflective of disease-causative expression alterations in these conditions. Detection of cell-specific transcriptional changes can also help develop more accurate disease models harboring these cellular alterations. Further, discovery of cell-specific transcriptional alterations in disease may uncover expression changes, particularly in less abundant cell types, which may be missed by the analysis of bulk transcriptome. Thus, there is a growing effort to identify cell-specific expression alterations in human diseases^9-11,14,15,17,46-48^.

There are two general approaches to decipher cell-specific transcriptional changes in AD. One approach is to conduct single nucleus (snRNAseq), single cell (scRNAseq) or purified cell RNAseq experimentally, followed by data analyses. The alternative approach is to design relatively complex bioinformatics pipelines to decipher cell-intrinsic information of individual cell types from bulk tissue microarray or RNAseq data. The first approach is limited due to significantly higher cost and experimental challenges. Additionally, these approaches may have the drawback that the procedures of dissociating cells break cell-to-cell communication and thus may not reflect the true expression signal *in vivo*. The alternative bioinformatics approach to decipher cell-specific transcriptome alterations from bulk tissue has the potential to avoid the above weaknesses of the experimental approach. Furthermore, a bioinformatics approach can make use of the large amount of existing RNAseq data^4-8,12^ from fresh or frozen bulk brain tissues with minimum added cost, and may better reflect the true situation in brain where different cells are in communication.

In this study, applying three different algorithms^13,14^ including our own novel approach, we estimated cell-intrinsic gene expression for deconvoluted cell types from three large bulk RNAseq datasets^4,8,12,18^ from two brain regions. We identified consensus cell-intrinsic transcriptional alterations (CI-DEGs) in AD, which are conserved across cohorts and brain regions. We also performed an in-depth comparison of these CI-DEGs with bulk brain RNAseq data obtained from the same datasets, collectively comprised of 685 unique brain samples. To our knowledge, this is the first study to detect CI-DEGs and their enriched gene ontology (GO) terms in computationally deconvoluted large-scale RNAseq data from AD and control brain samples. Additionally, we conducted a detailed comparison of CI-DEGs deconvoluted from bulk-DLPFC data with the three algorithms and those obtained from snRNAseq^19^ (snDLPFC) of a subset of the samples from the same cohort^7,8^.

The main findings from our study can be summarized as follows: **1)** The direction of change in cellular proportions in AD is consistent across two brain regions and three datasets for most cell types, although the magnitude of change seems to vary. Our findings revealed greater neuronal loss and microgliosis in TCX compared to DLPFC. **2)** We identified CI-DEGs and bulk tissue DEGs (bulk-DEGs) independently in two TCX and one DLPFC datasets. The overlap in bulk-DEGs across datasets is greater than that for CI-DEGs. **3)** We performed an in-depth comparison of the consensus CI-DEGs, common to both TCX and DLPFC against the consensus bulk-DEGs detected in these same datasets. We identified significant overlap between consensus CI-DEGs and consensus bulk-DEGs. The extent of overlap between consensus bulk-DEGs and consensus CI-DEGs was greatest for *down-regulated neuronal* genes (p=9.7E-206). This was followed by *up-regulated non-neuronal* genes (p ranging from 1.6E-77 for endothelia to 1.1E-17 for microglia). **4)** Despite the statistically significant overlap between consensus bulk vs. CI-DEGs, the majority of the consensus CI-DEGs for *up-regulated neuronal* and *down-regulated non-neuronal* genes were not detected in bulk tissue. This finding highlights the potential ability of computational deconvolution approach to identify CI-DEGs that may be missed in bulk-DEGs especially for genes that are not moving in the direction of cell proportion changes. **5**) We identified GO-terms enriched for consensus CI-DEGs, and detected processes that have previously been implicated in AD as well as novel ones. **6**) Using an snRNAseq^19^ dataset as comparison, we assessed the performance of our CI-DEG detection algorithm (WLC), and the published CellCODE^14^ and PSEA^13^ approaches. We determined that WLC had comparable or superior performance in the detection CI-DEGs that had significant overlap with snDLPFC results.

Our findings highlight the consistency and reproducibility of our findings across two different brain regions from three different studies conducted separately. We identified similar directions of change in AD:Control cell proportions in TCX and DLPFC. As expected from known AD neuropathology, neuronal populations are significantly lower, and microglial populations are significantly higher in AD vs. control brains in all datasets. Consistent with this pattern of reproducibility, we also found significant overlap of consensus CI-DEGs detected in TCX and DLPFC for all cell types and for both directions of change, i.e. up or down in AD, with consensus bulk-DEGs.

Using consensus CI-DEGs, we identified GO terms, which include processes and genes that have previously been implicated in AD, thereby providing further validation of our approach. Detailed discussion of all of the pathways identified with the consensus CI-DEGs is beyond the scope of this study. Instead, we herein highlight some of the CI-DEG enriched pathways.

Down-regulated neuronal CI-DEGs were enriched in neuropeptide hormone activity (GO:0005184) pathway. These terms include *VGF* (a.k.a. neuroendocrine regulatory peptide 1), which is selectively expressed in some neurons and shown to be reduced in AD and Parkinson’s disease^22^. Corticotropin releasing hormone (CRH), which is a neuronally expressed peptide that mediates stress in the hypothalamic-pituitary-adrenal axis, is also a member of the same GO term. CRH has been implicated in both adverse outcomes related to AD pathology in model systems and epidemiology studies, as well as having an important role in learning and memory^23^. Neuronal reduction of *CRH* may either be a potentially neuro-protective response in AD brains or lead to further negative impact on memory. Although the biological implications of our finding remain to be uncovered, our results are aligned with the potential importance of the neuroendocrine system in AD.

Interestingly, CI-DEGs also implicate biological processes that are enriched for neuronal-DEGs that are *<u>up</u>* in AD, despite reductions in neuronal cell populations in this condition for both TCX and DLPFC. This suggests that our cell type specific transcriptome deconvolution successfully captures transcript changes that occur in the direction opposite to that of cell-population changes, and that may therefore be missed in bulk-DEG approaches.

Indeed, the significant GO terms potassium channel activity (GO:0005267) and regulation of ion transport (GO:0043269) harbor many potassium channels, which are up in AD for neuronal CI-DEGs in both TCX and DLPFC, but do not have consistent results in bulk-DEGs from the same cohorts. These findings highlight the potential utility of cell-specific transcript deconvolution approaches in reducing noise from cell population changes, thereby resulting in consistent detection of true signal. Notably, potassium channels have been reported to be up in AD brains and mouse models of AD^49-51^, leading to their suggestion as a potential therapeutic avenue in this condition.

Notably, the computational approach we describe can be extended to cellular sub-types. To exemplify this, we utilized published snRNAseq^19^ data to identify marker genes of excitatory and inhibitory neurons. Using those and the previously applied markers for the glial cells, we ran DSA algorithm [3] to estimate cell proportion (**Fig S6**). We applied PSEA, WLC and CellCODE to identify consensus CI-DEGs in the same fashion as described (**Table S20, Fig S7**). As expected, both excitatory and inhibitory neuron proportions are both significantly lower in AD samples. Interestingly, the up-regulated excitatory neuronal consensus CI-DEGs are enriched in cation channel activity, which is not observed in inhibitory neurons (**Tables S21-S23**). These results highlight the ability of this approach to refine transcriptional alterations from bulk brain data to cellular sub-types.

Some consensus CI-DEGs point to AD-related perturbations of key cellular functions for the specific cell type. An example of this is consensus oligodendrocyte CI-DEGs. The *down-regulated* oligodendrocyte CI-DEGs are enriched for the myelination GO term (GO:0042552) and those that are *up* in this cell type are enriched in ceramide biosynthetic process (GO:0046513).

Down-regulated oligodendrocyte CI-DEGs include myelin basic protein (*MBP*)^4^, plasmolipin (*PLLP*)^4,5^, myelin and lymphocyte protein (*MAL*), and myelin-associated glycoprotein (*MAG*)^26^, even though bulk-DEGs for these genes did not show consistent changes. We^4^ and others^5^ demonstrated lower levels in AD of co-expression networks of genes implicated in myelination, which is consistent with the present findings from oligodendrocyte CI-DEGs.

Ceramide dysregulation has been implicated in AD^52,53^. Increased ceramide species were observed in AD and other neuropathological disorders compared to controls^53^, and the activation of the neutral sphingomyelinase–ceramide pathway induces oligodendrocyte death^54^. Our present observation from oligodendrocyte CI-DEGs are consistent with these findings.

Another potential utility of the computational deconvolution approach is its complementarity to snRNAseq data. While the latter is able to provide transcriptional data at a cellular level, it can miss information from low abundance cell populations. Indeed, in a recent snRNAseq^19^ study of AD and control brain samples, only 121 endothelial cells were identified out of a total of 70,634 cells; and consequently no endothelial DEGs were reported. In contrast, our computational approach identified 140 genes that are up-regulated in endothelial cells in AD and 72 genes that are down-regulated. We further validated the endothelial expression of these genes by confirming their expression in sorted endothelial cells^9^. The significantly enriched GO terms include cytoskeleton organization and translational initiation, for these genes up- and down-regulated, respectively, in endothelia (**Fig 3, Tables S12, S17**). These findings highlight the ability of the computational deconvolution approach to provide biological insights that could be missed by snRNAseq approaches.

Despite the biological insights gained from computationally deconvoluted CI-DEGs, they also have some shortcomings. Compared to bulk-DEGs, CI-DEGs between different datasets show less degree of overlap, regardless of the deconvolution algorithm utilized. This highlights the challenge in the field for the ultimate goal of minimizing detection of transcriptional perturbations due to cell proportion changes while maximizing discovery of those that lead to disease. Put differently, CI-DEGs may enhance discovery of true cell-specific transcriptional changes but this may come at the expense of increased false negative findings. In contrast, bulk-DEGs may capture a larger number of perturbed transcripts, but some may be merely due to cell population differences between diseased and healthy tissue. Ultimately, findings from both approaches may provide the greatest utility in detecting true positives.

Another analytic caveat is that cell proportion estimation approaches, including those used in this study yield relative values rather than the absolute levels of cell proportions. Detection of absolute levels in a typical bulk RNAseq experiment will be challenging if not impossible due to different mRNA amounts per cell type and the library preparation protocols (e.g. TruSeq^®^ RNA Sample Preparation v2 Guide) which require similar amounts of cDNA from each sample to be sent to the sequencer. This leads to a normalization which would preclude the detection of the absolute cell proportions, as illustrated in an example in **Fig. S8**. Despite these challenges, our findings reveal that analytic deconvolution of bulk RNAseq can detect cell-specific transcriptional changes.

Comparison of CI-DEGs deconvoluted from bulk-DLPFC and those detected using an orthogonal approach of snRNAseq^19^ from the same cohort (ROSMAP) demonstrates the ability of these computational approaches to identify true cell-specific expression changes in AD. Using our in-house WLC algorithm, there was significant overlap with the results from snRNAseq and CI-DEGs of most cell types (**Fig 4, Tables S18-S19**). CellCODE and PSEA also identified significant overlaps, but to a lesser extent, especially for rarer cell types such as microglia. Hence, our WLC algorithm demonstrates at least comparable performance to these two algorithms^13,14^. This is also supported by the higher degree of overlap among different cohorts for CI-DEGs detected by WLC than the other two algorithms (**Fig 2a**). Due to the challenges in deconvoluting noisy data from human series, different computational approaches may be utilized and combined, and that calls for a more devoted effort in developing such algorithms.

In summary, using three distinct computational approaches, we deconvoluted brain bulk-RNAseq data from three large and independent cohorts^8,12,18^. We detected cell population changes that are observed consistently across cohorts, and congruent with the known disease pathology. Although there is significant overlap between consensus CI-DEGs and consensus bulk-DEGs, there are more unique CI-DEGs that change in the direction opposite to that of cell population changes. This suggests that CI-DEGs may have utility in detecting disease-related transcriptional changes above and beyond those due to cell proportion changes. Consensus CI-DEGs identify GO terms, including those for hormone activity, myelin biology and channel activity. The enriched CI-DEGs include genes previously implicated in AD or neurodegeneration, such as *VGF, CRH, MOBP* and *MBP*, and other novel genes.

## Methods

### Analysis datasets

We generated the TCX-Mayo data, which consists of temporal cortex RNAseq measurement of 80 AD patients and 28 controls diagnosed according to neuropathologic criteria^4,12^. RNAseq data were processed and quality control (QC) was conducted as described^4,12^. ROSMAP DLPFC^7,8^ and TCX-MSBB^18^ datasets were downloaded from the AMP-AD Knowledge Portal on Synapse (syn8691134 and syn8691099). We further filtered out non-Caucasian samples and those that had incongruent sex based on provided covariate vs. transcriptome data. All samples were classified as AD or control based on neuropathological data. All TCX-Mayo AD samples had Braak neurofibrillary tangle (NFT) score ≥4. TCX-Mayo controls had Braak score ≤3 and were without any neurodegenerative disease diagnoses. TCX-MSBB AD samples had Braak ≥4 and CERAD ≥2; and controls had Braak ≤3 and CERAD ≤1. DLPFC AD samples form ROSMAP had Braak score ≥4 and CERAD neuritic plaque score ≤2. ROSMAP control samples had Braak ≤3 and CERAD neuritic plaque score ≥3.

It should be noted that the CERAD^55^ neuritic plaque score as applied by the ROSMAP study is defined such that high CERAD indicates lower neuritic plaque burden and decreased probability of AD. In the MSBB study, higher CERAD indicates higher plaque burden.

Raw RNA read counts were normalized using conditional quantile normalization (CQN) method^56^ implemented in R cqn package, as previously described^4^. This normalization takes into consideration library size, gene length and GC content. It also performs a log2 transformation so that the resulting distribution for each gene is Gaussian-like. We also determined covariates that contributed significantly to the variation of gene expression in these RNAseq cohorts (**Fig S5**) for adjustment in the analyses.

### Cell proportion estimation

Digital sorting algorithm (DSA)^16^ was applied to estimate cell proportions through R DSA package, function DSA. For each cell type, DSA first computes the average marker gene expression in the analysis dataset, the purpose of which is to construct a variable that better reflects cell proportion variation among subjects. To reduce the effect of outlier expression that is occasionally seen in RNAseq data, we modified the original DSA so that the median instead of mean expression was computed.

### CI-DEG analysis for individual cell types

In this study we identified CI-DEGs from deconvoluted bulk RNAseq data using three different algorithms, namely PSEA^13^, CellCODE^14^ and our in-house algorithm WLC. All analyses adjusted for the following variables: Sex, RIN, age at death and batch for DLPFC and TCX-MSBB datasets, and sex, RIN and age at death for TCX-Mayo dataset (**Fig S5**).

PSEA^13^ applies model selection procedures to select cell type(s) that should be included in baseline (control) or AD condition, and then estimate differential expression in specific cell types (CI-DEGs). We used the R package PSEA to obtain CI-DEG results of PSEA approach, through functions em_quantvg (to generate candidate models) and lmfitst (to fit all candidate models and pick the best one). Expression values used in PSEA are in linear scale (non log-transformed).

CellCODE^14^ assesses overall gene expression difference between AD and control groups with adjustment of relative cell proportion, followed by assigning the cell type that correlates best with the difference (CI-DEGs). R package CellCODE was used to obtain DEG results of CellCODE approach, through functions getAllSPVs (to construct surrogate variable using marker genes through singular value decomposition) and lm.coef (to estimate difference between AD and control groups). Strictly speaking, CellCODE measures the overall differential expression rather than CI-DEGs but identifies the cell type that is most correlated with the observed difference between AD and control using cellPopT function. However, for simplicity, we refer to the DEGs from CellCODE as CI-DEGs in this study. Expression values used in CellCODE are in log scale.

Our in-house method WLC, described in the method section, applies <u>w</u>eighted linear regression with <u>c</u>onstraints and model selection procedures, which also estimates differential expression in specific cell types (CI-DEGs). It guarantees the fitted relative gene expression is non-negative. By weighing the expression, it smooths out the extreme data points. The procedures of this algorithm is illustrated by the following high level pseudo code.

> *Assume cell type 1,2,3,4,5 are neu, oli, mic, ast and end respectively*
>
> *Fit Eq.(1) to identify a set of cell type T in which the gene is significantly expressed If the set T is not empty:*
>
> *Fit Eq.(2) to identify a set of cell type Φ ⊆ T in which the gene is differentiallly expressed*
>
> *Let Ω = {each cell type in T, Φ, T}*
>
> *For each element Θ* ∈ *Ω*
>
> *Fit Eq.(3) with adjustment for other covariates C*_*k*_*(1≤ k ≤ m)*
>
> *Keep Akaike information criterion (AIC) of this model fitting*
>
> *Identify Θ*_*best*_ *that gives the best AIC.*
>
> *Use the estimated values from the best model and obtain p-value from F-test*

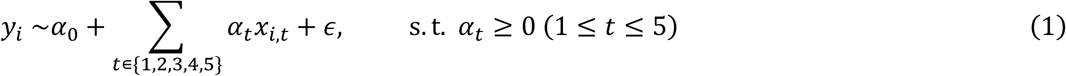

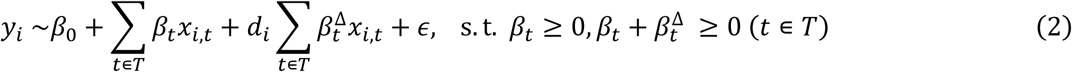

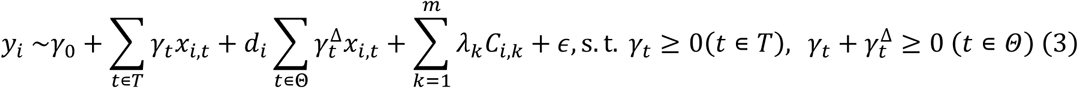

In Eqs.1-3, *y*_*i*_ is the observed expression of a gene in subject i; x_i,t_ is the median marker gene expression of cell type t in subject i; *C*_*i,k*_ is covarite *k* in subject i. In Eq.1, *α*_*t*_ is the overall relative expression in cell type t. In Eq.2, β_t_ is relative expression at the baseline condition in cell type t; d_i_ = 0 if subject i is in control group, and d_i_ = 1 if subject i is in AD group; therefore, 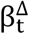 is the difference of relative gene expression between baseline condition and AD condition in cell type t. Of note, due to the biological meaning these coefficents, they must satisfy constraints such that *α*_*t*_, *β*_*t*_, 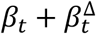 be non-negative. In addition, y_i_ is in linear scale rather than in log scale^57^, and these non-log-transformed expression values tend to have extreme data points that need to be weighted down. Based on the above considerations, Eq. 1 was fitted by weighted least square linear regression with contraints, which is implemented in lsei function in R package limSolve. The weight of each observation (*w*_*i*_) is determined by formulae Eq.3 and Eq.4 below. Intuitively, if the expression of a gene in a sample is extremely distant from the median expression of all samples in the same diagnosis group, the weight of that sample is smaller than 1 for that gene.

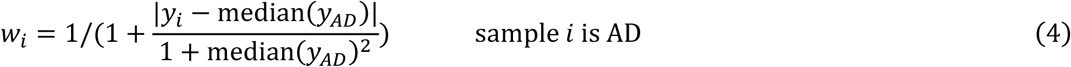

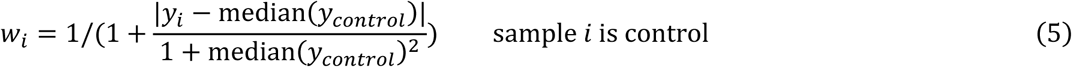

### GO enrichment analysis

Using genes included in the CI-DEG analysis as background genes, p-value for GO term enrichment with consensus CI-DEGs was calculated by “enrichmentAnalysis” function from WGCNA R package^35^.

### Comparison of CI-DEGs from computational deconvolution vs. snRNAseq

To determine if computational bulk-tissue RNAseq could reveal true CI-DEGs, we downloaded and utilized a published snRNAseq study^19^ from frozen DLPFC tissues (snDLPFC) which compared gene expression from 24 individuals with AD-pathology to that from 24 individuals without AD-pathology in six cell types - excitatory neurons, inhibitory neurons, oligodendrocyte, microglia, oligodendrocyte precursor cells and astrocytes. These snDLPFC samples are from the ROSMAP project, of which we analyzed 474 bulk-DLPFC RNAseq data in our current study. Twenty-four (9 AD cases, 15 controls) of the 48 individuals in snDLPFC study are also included in the bulk-DLPFC dataset. Hence both the snDLPFC and bulk-DLPFC are from the same cohort with significant overlap. Three deconvolution methods were included in this comparison – CellCODE^14^, PSEA^13^ and WLC, our in-house method.

Two types of comparisons were made between the deconvoluted bulk-DLPFC and snDLPFC results. In the first comparison, we ranked the genes by their p-values of differential expression between AD and control subjects, per cell type. We compared the top *N* up- or down-genes from the snDLPFC study to those identified by each deconvolution algorithm, per cell type. Genes common to both the deconvoluted bulk-DLPFC and snDLPFC were counted and plotted.

In the second set of comparisons, CI-DEGs identified from deconvolution methods at nominal significance (p-value <0.05) were compared to those identified in the snRNAseq data (p-value <0.05).

To assess if the observed overlap from each set of analyses is significant with regards to overlap between random selections, we obtained empirical p values from computer simulations described below.

### Empirical p-value for the number of overlapping genes

The empirical p-values for the number of overlapping genes between snDLPFC and bulk-DLPFC was obtained using a computer simulation as follows. (**A**) Let *S*_*sn*_ stand for all genes in snDLFC and *S*_*bulk*_ for all genes in bulk-DLFC; (**B**) randomly assign up-regulation on each *S*_*sn*_ gene at probability 0.5, and on each *S*_*bulk*_ gene at probability 0.5; (**C**) randomly pick *N* genes from up-genes of *S*_*sn*_, randomly pick *N* genes from up-genes of *S*_*bulk*_, and count the number of overlapping genes; (**D**) Steps B-C were repeated 10000 times, and the numbers of overlaps (*M*_1_, *M*_2_, …, *M*_10000_) were obtained; (**E**) Let *M* be the number of observed overlapping genes, and the empirical p-value is (1+number of occurrences that *M*_*i*_ >= *M*)/10001.

### Identification of excitatory and inhibitory neuronal markers

Using a published human brain snRNAseq dataset ^19^, we identified excitatory and inhibitory neuronal markers using Seurat R package FindMarker function. Excitatory neuronal markers are those that **a)** are detected in >= 70% of excitatory neurons, **b)** have average normalized expression in excitatory neurons that are >= 4.5X of that in each of the other cell types (i.e. inhibitory neurons, oligodendrocytes, microglia, astrocytes and OPC), and **c)** have rank sum test p-value < 0.05 in the comparison of expression levels in excitatory neurons and each of the other cell types. Inhibitory neuronal markers were similarly identified except that we required a less stringent detection limit of >= 50% of inhibitory neurons as the >= 70% threshold yielded too few genes. Excitatory neuronal markers were further refined by requiring their presence in the neuronal markers from BRETIGEA ^15^. A total of 53 excitatory and 62 inhibitory neuronal markers were identified (**Table S20**). Excitatory and inhibitory CI-DEGs were identified as described above.

## Conclusions

This study demonstrates the utility of our analytic approach in deciphering cell-specific transcriptional alterations using bulk tissue in a complex disease, provides a comprehensive comparison of our pipeline to existing ones, identifies patterns of cell proportions in AD and control samples across brain regions, discovers novel CI-DEGs with replication across independent cohorts and highlights biological processes with cell-specific expression changes in AD. These findings are expected to refine discovery of molecular therapeutic targets, biomarkers that reflect cellular transcriptional alterations in AD and accelerate generation of more accurate disease models.

## Abbreviations

AD: Alzheimer’s disease
ADCY8: Adenylate cyclase 8
AMP-AD: Accelerating Medicines Partnership Alzheimer’s Disease
Bulk-DEGs: Differentially expressed genes from bulk brain tissue
Bulk-DLPFC: Bulk RNA sequencing from dorsolateral prefrontal cortex
C3AR1: Complement C3a receptor 1
CEBPA: CCAAT enhancer binding protein alpha
CERS4: Ceramide synthase 4
CI-DEGs: Cell-intrinsic differentially expressed gene(s)
CNS: Central nervous system
CQN: Conditional quantile normalization
CRH: corticotropin releasing hormone
DEGs: Differentially expressed genes
DLPFC: Dorsolateral prefrontal cortex
DSA: Digital sorting algorithm
GJA1: Connexin43
GO: Gene Ontology
GRIN1: Glutamate ionotropic receptor NMDA type subunit 1
GSTM2: Glutathione S-transferase mu 2
IL18: Interleukin 18
MAG: Myelin-associated glycoprotein
MAL: Myelin and lymphocyte protein
MBP: Myelin basic protein
NFT: Neurofibrillary tangle
OPC: Oligodendrocyte progenitor cell
PLLP: Plasmolipin
QC: Quality control
RIN: RNA integrity number
RNAseq: RNA sequencing
ROSMAP: Rush Religious Orders Study and Memory and Aging Project
scRNAseq: Single cell RNA sequencing
snDLPFC: Single nucleus RNA sequencing from dorsolateral prefrontal cortex
snRNAseq: Single nucleus RNA sequencing
SYT1: Synaptotagmin
TCX: Temporal cortex
TCX-Mayo: Mayo Clinic temporal cortex
TCX-MSBB: Mount Sinai VA Medical Center Brain Bank temporal cortex
TXN2: Thioredoxin2
UGT8: UDP glycosyltransferase 8
VGF: VGF Nerve Growth Factor Inducible, a.k.a neuroendocrine regulatory peptide 1
WGCNA: Weighted gene co-expression network analysis

## Declarations

### Ethical Approval and Consent to participate

All data were generated from deceased individual’s autopsied brain tissue. This study was approved by Mayo Clinic Institutional Review Board.

### Consent for publication

All authors reviewed and approved the final manuscript.

### Availability of supporting data

The data used in this manuscript is available to the research community through the AMP-AD knowledge portal on Sage Synapse as follows:

| Dataset | Data Type | Description | SynapseID | DoD |
| --- | --- | --- | --- | --- |
| Mayo RNAseq TCX | RNASeq Expression | MAPRseq processed, CQN normalized counts | * | na |
| Mayo RNAseq TCX | Metadata | Individual human and RNAseq | syn3817650 | na |
| Mayo RNAseq TCX | Metadata | Quality Control | syn6126114 | na |
| ROSMAP | RNASeq Expression | Consensus processed RNASeq raw counts | syn8691134 | 10/2/2019 |
| ROSMAP | Metadata | ID Key | syn3382527 | 10/2/2019 |
| ROSMAP | Metadata | Individual human | syn3191087 | 10/2/2019 |
| ROSMAP | Metadata | Assay<br>RNAseq | syn21088596 | 1/2/2020 |
| MSBB | RNASeq<br>Expression | Consensus<br>processed<br>RNASeq<br>raw counts | syn8691099 | 10/2/2019 |
| MSBB | Metadata | Individual<br>human | syn6101474 | 11/22/2019 |
| MSBB | Metadata | Assay<br>RNAseq | syn6100548 | 10/2/2019 |
| snRNAseqPFC_BA10 | Gene<br>Expression | Count matrix | syn18686381 | 11/1/2019 |
| snRNAseqPFC_BA10 | Metadata | Column<br>metadata | syn18686372 | 11/1/2019 |
| snRNAseqPFC_BA10 | Metadata | Gene row<br>names | syn18686382 | 11/1/2019 |
| snRNAseqPFC_BA10 | Metadata | Biospecimen<br>metadata | syn18642936 | 11/1/2019 |

DoD = Date of download, “na” indicates data that was generated by study authors and shared within the AMP-AD knowledge portal. *Data not yet available on the AMP-AD knowledge portal, but will be uploaded and shared upon acceptance for publication. SynapseIDs can be searched directly within the portal only (https://adknowledgeportal.synapse.org/). Scripts to perform the analysis and results reported here will be made available upon acceptance of the manuscript for publication.

### Author contributions

X.W., S.L., M.A. and N.E-T conceived the idea. X.W., M.A. and N.E-T developed and designed the study, and wrote the manuscript. X.W. and S.L. developed the mathematical theory and X.W. performed the computations.D.D. provided tissue from the Mayo Clinic Brain Bank and neuropathologically characterized TCX-Mayo brain samples. T.N., K.M., S.L., M.A. and M.C. performed sample selection and library preparation of the TCX-Mayo dataset. M.A., Z.Q., T.C., T.P. and J.R. performed quality control of DLPFC and TCX-MSBB datasets. J.C. and Y.A. supervised the analytical aspects of the project. N.E-T. provided funding, supervision and direction for the whole study. All authors read and approved the final manuscript.

## Acknowledgements

We thank the patients and their families for their participation, without them these studies would not have been possible. The Mayo RNAseq study data was led by Dr. Nilüfer Ertekin-Taner, Mayo Clinic, Jacksonville, FL as part of the multi-PI U01 AG046139 (MPIs Golde, Ertekin-Taner, Younkin, Price) using samples from The Mayo Clinic Brain Bank. Data collection was supported through funding by NIA grants P50 AG016574, R01 AG032990, U01 AG046139, R01 AG018023, U01 AG006576, U01 AG006786, R01 AG025711, R01 AG017216, R01 AG003949, NINDS grant R01 NS080820, CurePSP Foundation, and support from Mayo Foundation. This study was in part funded by NIH RF1 AG051504 and R01 AG061796 (NET). We thank the Mayo Clinic Advanced Genomic Technology Center staff and bioinformatics core facility for all gene expression measurements. XW thanks Dr. E. Aubrey Thompson for text editing and Jeremy Burgess for provide insightful inputs. XW was funded in part by a pilot grant from the Mayo Clinic ADRC (P50 AG016574).

The results published here are in whole or in part based on data obtained from the AMP-AD Knowledge Portal (https://adknowledgeportal.synapse.org). Study data were provided by the Rush Alzheimer’s Disease Center, Rush University Medical Center, Chicago. Data collection was supported through funding by NIA grants P30AG10161 (ROS), R01AG15819 (ROSMAP; genomics and RNAseq), R01AG17917 (MAP), R01AG30146, R01AG36042 (5hC methylation, ATACseq), RC2AG036547 (H3K9Ac), R01AG36836 (RNAseq), R01AG48015 (monocyte RNAseq) RF1AG57473 (single nucleus RNAseq), U01AG32984 (genomic and whole exome sequencing), U01AG46152 (ROSMAP AMP-AD, targeted proteomics), U01AG46161(TMT proteomics), U01AG61356 (whole genome sequencing, targeted proteomics, ROSMAP AMP-AD), the Illinois Department of Public Health (ROSMAP), and the Translational Genomics Research Institute (genomic).

The results published here are in whole or in part based on data obtained from the AMP-AD Knowledge Portal (https://adknowledgeportal.synapse.org/). These data were generated from postmortem brain tissue collected through the Mount Sinai VA Medical Center Brain Bank and were provided by Dr. Eric Schadt from Mount Sinai School of Medicine.

## Author’s information

Not applicable.

## References

1. 2018 Alzheimer’s disease facts and figures. Alzheimer’s & Dementia 14, 367–429 (2018).

2. Mehta, D., Jackson, R., Paul, G., Shi, J. & Sabbagh, M. Why do trials for Alzheimer’s disease drugs keep failing? A discontinued drug perspective for 2010-2015. Expert opinion on investigational drugs 26, 735–739 (2017).

3. Anderson, R.M., Hadjichrysanthou, C., Evans, S. & Wong, M.M. Why do so many clinical trials of therapies for Alzheimer’s disease fail? The Lancet 390, 2327–2329 (2017).

4. Allen, M. et al. Conserved brain myelination networks are altered in Alzheimer’s and other neurodegenerative diseases. Alzheimer’s & Dementia (2017).

5. McKenzie, A.T. et al. Multiscale network modeling of oligodendrocytes reveals molecular components of myelin dysregulation in Alzheimer’s disease. Molecular Neurodegeneration 12, 82 (2017).

6. Zhang, B. et al. Integrated systems approach identifies genetic nodes and networks in late-onset Alzheimer’s disease. Cell 153, 707–720 (2013).

7. Mostafavi, S. et al. A molecular network of the aging human brain provides insights into the pathology and cognitive decline of Alzheimer’s disease. Nature Neuroscience 21, 811–819 (2018).

8. De Jager, P.L. et al. A multi-omic atlas of the human frontal cortex for aging and Alzheimer’s disease research. Scientific Data 5, 180142 (2018).

9. Zhang, Y. et al. Purification and Characterization of Progenitor and Mature Human Astrocytes Reveals Transcriptional and Functional Differences with Mouse. Neuron 89, 37–53 (2016).

10. Darmanis, S. et al. A survey of human brain transcriptome diversity at the single cell level. Proceedings of the National Academy of Sciences 112, 7285–7290 (2015).

11. Lake, B.B. et al. Integrative single-cell analysis of transcriptional and epigenetic states in the human adult brain. Nature biotechnology 36, 70–80 (2018).

12. Allen, M. et al. Human whole genome genotype and transcriptome data for Alzheimer’s and other neurodegenerative diseases. Scientific Data 3, 160089 (2016).

13. Kuhn, A., Thu, D., Waldvogel, H.J., Faull, R.L.M. & Luthi-Carter, R. Population-specific expression analysis (PSEA) reveals molecular changes in diseased brain. Nat Meth 8, 945–947 (2011).

14. Chikina, M., Zaslavsky, E. & Sealfon, S.C. CellCODE: a robust latent variable approach to differential expression analysis for heterogeneous cell populations. Bioinformatics 31, 1584–1591 (2015).

15. McKenzie, A.T. et al. Brain Cell Type Specific Gene Expression and Co-expression Network Architectures. Scientific Reports 8, 8868 (2018).

16. Zhong, Y., Wan, Y.-W., Pang, K., Chow, L.M. & Liu, Z. Digital sorting of complex tissues for cell type-specific gene expression profiles. BMC Bioinformatics 14, 89 (2013).

17. Kuhn, A. et al. Cell population-specific expression analysis of human cerebellum. BMC genomics 13, 610 (2012).

18. Wang, M. et al. The Mount Sinai cohort of large-scale genomic, transcriptomic and proteomic data in Alzheimer’s disease. Sci Data 5, 180185 (2018).

19. Mathys, H. et al. Single-cell transcriptomic analysis of Alzheimer’s disease. Nature (2019).

20. Ashburner, M. et al. Gene ontology: tool for the unification of biology. The Gene Ontology Consortium. Nature genetics 25, 25–29 (2000).

21. Langfelder, P. & Horvath, S. WGCNA: an R package for weighted correlation network analysis. BMC Bioinformatics 9, 559 (2008).

22. Cocco, C. et al. Distribution of VGF peptides in the human cortex and their selective changes in Parkinson’s and Alzheimer’s diseases. Journal of anatomy 217, 683–93 (2010).

23. Futch, H.S., Croft, C.L., Truong, V.Q., Krause, E.G. & Golde, T.E. Targeting psychologic stress signaling pathways in Alzheimer’s disease. Molecular neurodegeneration 12, 49 (2017).

24. Wang, R. & Reddy, P.H. Role of Glutamate and NMDA Receptors in Alzheimer’s Disease. J Alzheimers Dis 57, 1041–1048 (2017).

25. Berchtold, N.C. et al. Brain gene expression patterns differentiate mild cognitive impairment from normal aged and Alzheimer’s disease. Neurobiol Aging 35, 1961–72 (2014).

26. McAleese, K.E. et al. Parietal white matter lesions in Alzheimer’s disease are associated with cortical neurodegenerative pathology, but not with small vessel disease. Acta neuropathologica 134, 459–473 (2017).

27. Crivelli, S.M. et al. Sphingolipids in Alzheimer’s disease, how can we target them? Adv Drug Deliv Rev (2020).

28. Olsen, A.S.B. & Faergeman, N.J. Sphingolipids: membrane microdomains in brain development, function and neurological diseases. Open Biol 7(2017).

29. El Gaamouch, F. et al. VGF-derived peptide TLQP-21 modulates microglial function through C3aR1 signaling pathways and reduces neuropathology in 5xFAD mice. Molecular neurodegeneration 15, 4–4 (2020).

30. Tzeng, T.-C. et al. Inflammasome-derived cytokine IL18 suppresses amyloid-induced seizures in Alzheimer-prone mice. Proceedings of the National Academy of Sciences of the United States of America 115, 9002–9007 (2018).

31. White, C.S., Lawrence, C.B., Brough, D. & Rivers-Auty, J. Inflammasomes as therapeutic targets for Alzheimer’s disease. Brain Pathol 27, 223–234 (2017).

32. Gao, T. et al. Transcriptional regulation of homeostatic and disease-associated-microglial genes by IRF1, LXRβ, and CEBPα. Glia 67, 1958–1975 (2019).

33. Deitmer, J.W., Theparambil, S.M., Ruminot, I., Noor, S.I. & Becker, H.M. Energy Dynamics in the Brain: Contributions of Astrocytes to Metabolism and pH Homeostasis. Frontiers in Neuroscience 13(2019).

34. Bélanger, M., Allaman, I. & Magistretti, Pierre J. Brain Energy Metabolism: Focus on Astrocyte-Neuron Metabolic Cooperation. Cell Metabolism 14, 724–738 (2011).

35. Zhang, Q. et al. Integrated proteomics and network analysis identifies protein hubs and network alterations in Alzheimer’s disease. Acta neuropathologica communications 6, 19–19 (2018).

36. Lovell, M.A., Xie, C., Gabbita, S.P. & Markesbery, W.R. Decreased thioredoxin and increased thioredoxin reductase levels in alzheimer’s disease brain. Free Radical Biology and Medicine 28, 418–427 (2000).

37. Kajiwara, Y. et al. GJA1 (connexin43) is a key regulator of Alzheimer’s disease pathogenesis. Acta Neuropathol Commun 6, 144 (2018).

38. de Quervain, D.J. & Papassotiropoulos, A. Identification of a genetic cluster influencing memory performance and hippocampal activity in humans. Proc Natl Acad Sci U S A 103, 4270–4 (2006).

39. Shi, Y. et al. Rapid endothelial cytoskeletal reorganization enables early blood–brain barrier disruption and long-term ischaemic reperfusion brain injury. Nature Communications 7, 10523 (2016).

40. Stamatovic, S.M., Keep, R.F. & Andjelkovic, A.V. Brain endothelial cell-cell junctions: how to “open” the blood brain barrier. Current neuropharmacology 6, 179–192 (2008).

41. Evans, H.T., Benetatos, J., van Roijen, M., Bodea, L.G. & Gotz, J. Decreased synthesis of ribosomal proteins in tauopathy revealed by non-canonical amino acid labelling. EMBO J 38, e101174 (2019).

42. Garcia-Esparcia, P. et al. Altered machinery of protein synthesis is region- and stage-dependent and is associated with alpha-synuclein oligomers in Parkinson’s disease. Acta Neuropathol Commun 3, 76 (2015).

43. Koren, S.A. et al. Tau drives translational selectivity by interacting with ribosomal proteins. Acta Neuropathol 137, 571–583 (2019).

44. Conway, O.J. et al. ABI3 and PLCG2 missense variants as risk factors for neurodegenerative diseases in Caucasians and African Americans. Molecular neurodegeneration 13, 53 (2018).

45. Srinivasan, K. et al. Untangling the brain’s neuroinflammatory and neurodegenerative transcriptional responses. Nature communications 7, 11295 (2016).

46. Lin, M.-Y. et al. Releasing Syntaphilin Removes Stressed Mitochondria from Axons Independent of Mitophagy under Pathophysiological Conditions. Neuron 94, 595-610.e6 (2017).

47. Newberg, L.A., Chen, X., Kodira, C.D. & Zavodszky, M.I. Computational de novo discovery of distinguishing genes for biological processes and cell types in complex tissues. PloS one 13, e0193067 (2018).

48. Li, Z. et al. Genetic variants associated with Alzheimer’s disease confer different cerebral cortex cell-type population structure. Genome medicine 10, 43 (2018).

49. Angulo, E. et al. Up-regulation of the Kv3.4 potassium channel subunit in early stages of Alzheimer’s disease. J Neurochem 91, 547–57 (2004).

50. Maezawa, I. et al. Kv1.3 inhibition as a potential microglia-targeted therapy for Alzheimer’s disease: preclinical proof of concept. Brain 141, 596–612 (2018).

51. Yi, M. et al. KCa3.1 constitutes a pharmacological target for astrogliosis associated with Alzheimer’s disease. Mol Cell Neurosci 76, 21–32 (2016).

52. Czubowicz, K., Jęśko, H., Wencel, P., Lukiw, W.J. & Strosznajder, R.P. The Role of Ceramide and Sphingosine-1-Phosphate in Alzheimer’s Disease and Other Neurodegenerative Disorders. Molecular neurobiology 56, 5436–5455 (2019).

53. Filippov, V. et al. Increased ceramide in brains with Alzheimer’s and other neurodegenerative diseases. Journal of Alzheimer’s disease : JAD 29, 537–547 (2012).

54. Lee, J.-T. et al. Amyloid-beta peptide induces oligodendrocyte death by activating the neutral sphingomyelinase-ceramide pathway. The Journal of cell biology 164, 123–131 (2004).

55. Mirra, S.S. et al. The Consortium to Establish a Registry for Alzheimer’s Disease (CERAD). Part II. Standardization of the neuropathologic assessment of Alzheimer’s disease. Neurology 41, 479–86 (1991).

56. Hansen, K.D., Irizarry, R.A. & Wu, Z. Removing technical variability in RNA-seq data using conditional quantile normalization. Biostatistics 13, 204–216 (2012).

57. Zhong, Y. & Liu, Z. Gene expression deconvolution in linear space. Nat Meth 9, 8–9 (2012).

