## supplemental Figure for "Deciphering cellular transcriptional alterations in Alzheimer’s disease brains"

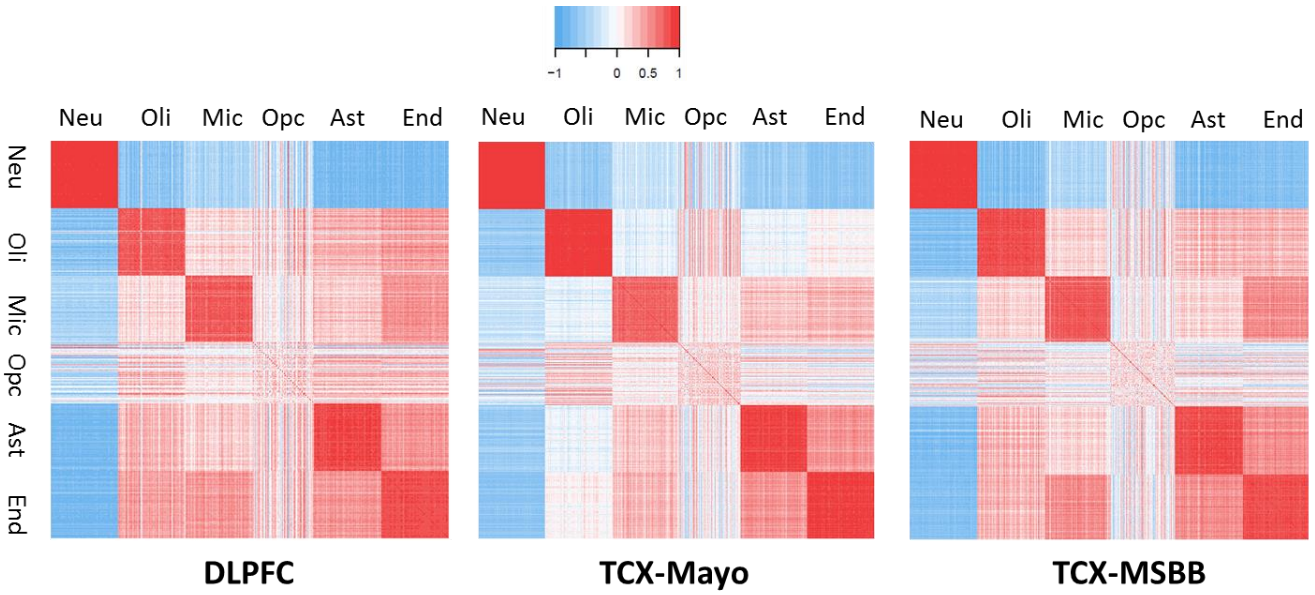


**Figure S1:** Pearson correlation of estimated cell proportions using different sets of marker genes randomly selected. The steps of this simulation study are as follows. **(1)** Obtain candidate markers from R package BRETIGEA[1] for neurons (top 500 out of 1000 listed), oligodendrocyte (top 500 out of 1000 listed), microglia (top 500 out of 1000 listed), OPC (top 250 out of 500 listed), astrocyte (top 500 out of 1000 listed) and endothelial (top 500 out of 1000 listed). **(2)** From candidate markers, randomly select 1/10 genes for each cell type, that is 50 selected genes for all cell types except OPC which has 25 selected genes. **(3)** Estimate cell proportion using DSA algorithm[2]. **(4)** Repeat (2)-(3) 100 times, and compute Pearson correlation of estimated cell proportion between different runs.

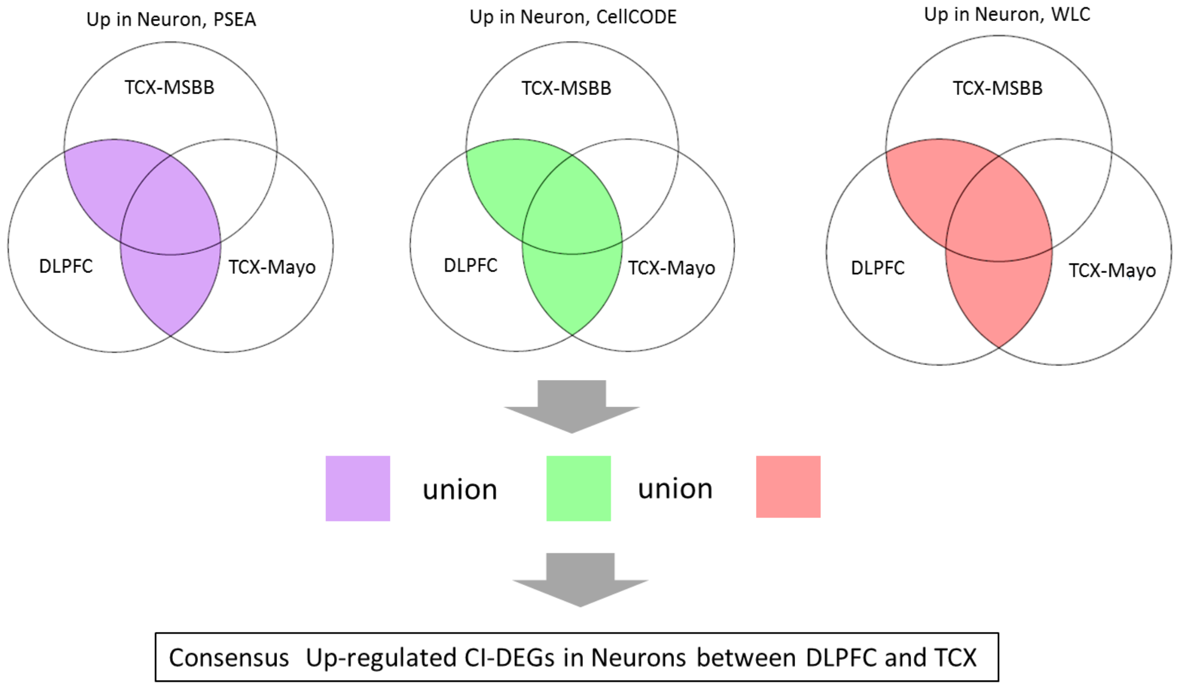


**Figure S2:** Illustration of obtaining consensus CI-DEGs between DLPFC and TCX regions, using up-regulated CI-DEGs in neuronal cells as an example.

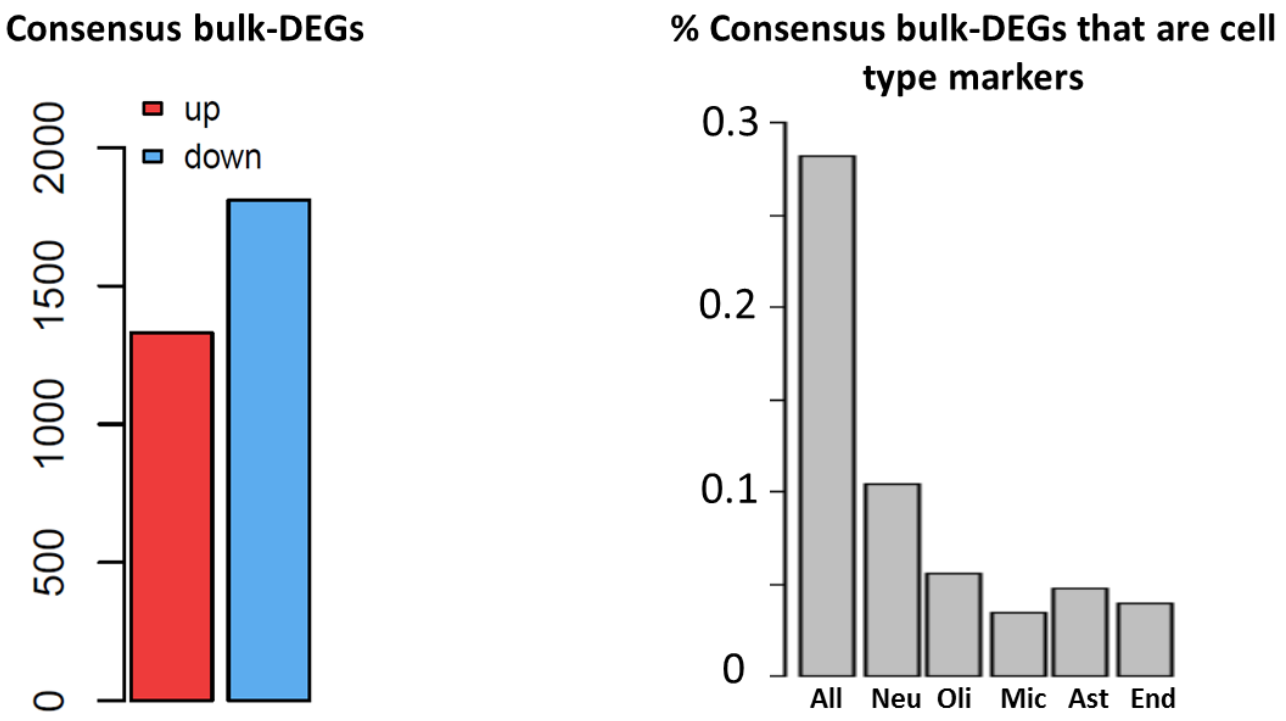


**Figure S3:** Left panel: the number of up-regulated and down-regulated consensus bulk-DEGs between DLPFC and TCX-Mayo, or between DLPFC and TCX-MSSM. Right panel: percent of consensus CI-DEGs that are also cell type marker genes. Cell type markers are from BRETIGEA[1], containing 1000 markers for each of the five cell types.

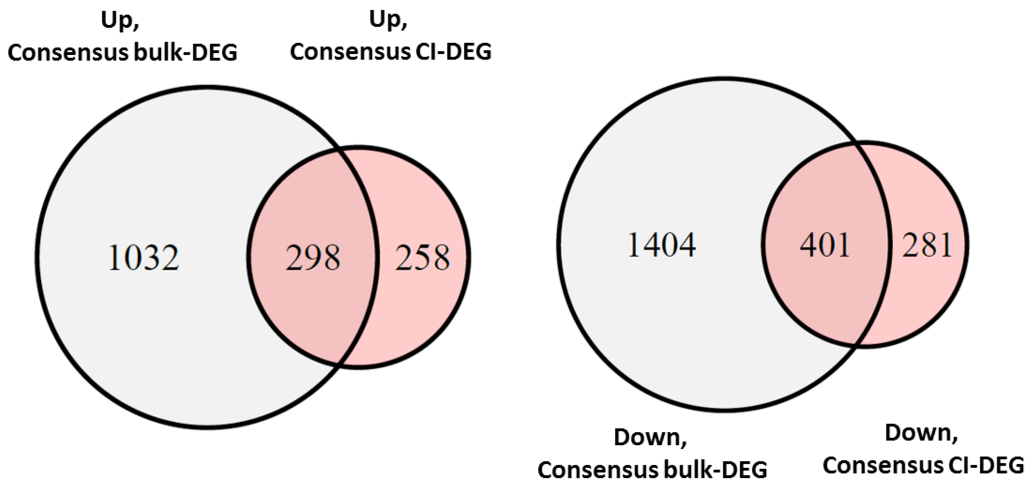


**Figure S4:** Venn diagram for all consensus bulk-DEGs and consensus CI-DEGs.

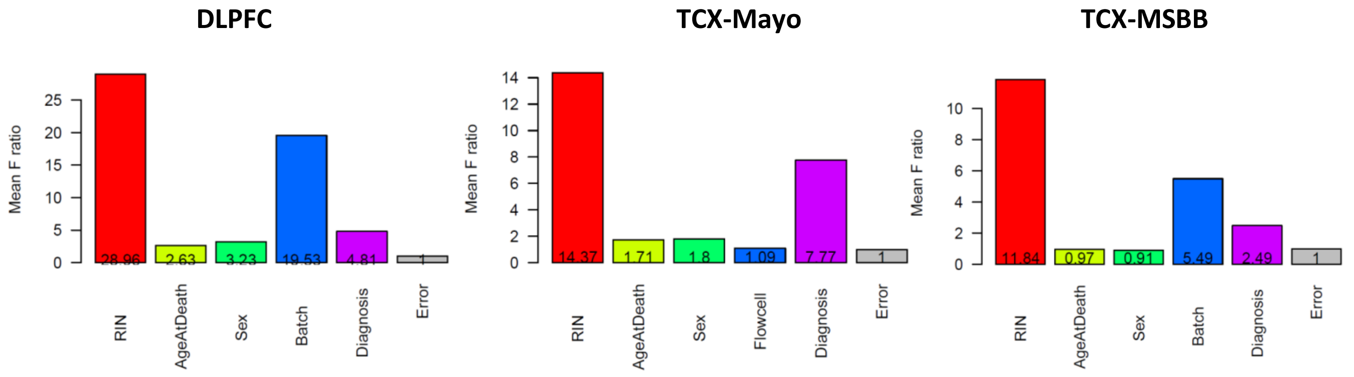


**Figure S5:** Source of variance analysis of in the DLPFC RNAseq dataset (left panel), TCX-Mayo (middle panel) and TCX-MSBB (right panel). For each gene, a full model was fitted in which cqn normalized gene expression with gene expression as dependent variable and RIN, age at death, sex, batch, and diagnosis group as independent variables (for DLPFC); RIN, age at death, sex, flowcell, and diagnosis group as independent variables (for TCX-Mayo); RIN, age at death, sex, batch, and diagnosis group as independent variables (for TCX-MSBB). Partial models were fitted using the same dependent variable and all but one independent variable. F statistics were obtained by comparing the full model and partial model for each independent variable. Y-axis is the mean of values of F statistics over all genes. In DLPFC and TCX-MSBB, diagnosis, age at death, sex, RIN and batch contributed more than random errors to the variation of gene expression, whereas in TCX-Mayo diagnosis, age at death, sex, and RIN contributed more than random errors to the variation of gene expression.

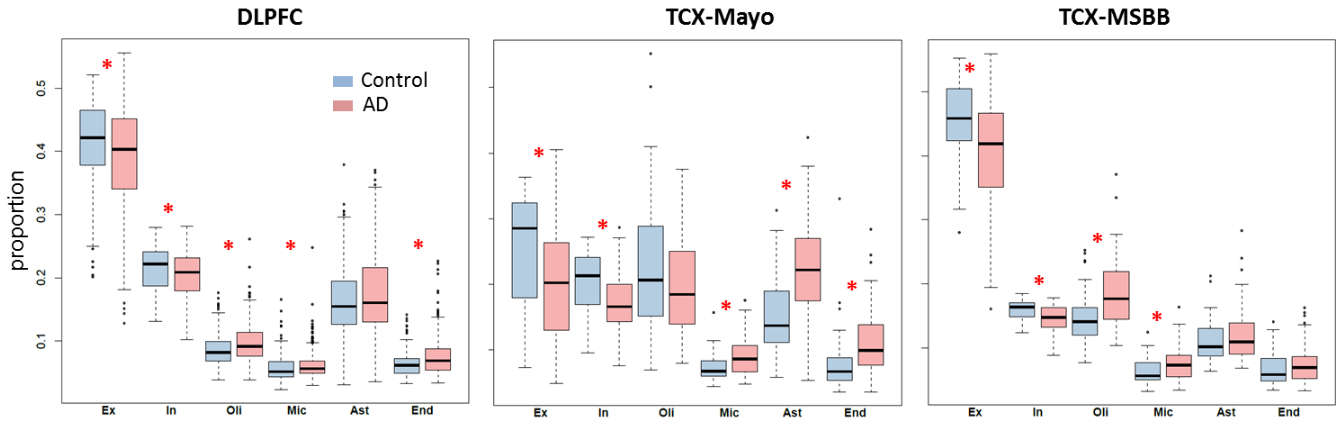


**Figure S6:** Estimated cell proportion of excitatory neuron, inhibitory neuron, oligodendrocyte, microglia, astrocyte and endothelial cells. Red asterisk indicates location shifts between cell proportions in AD and control groups at nominal p value 0.05 from Wilcoxon rank sum test.

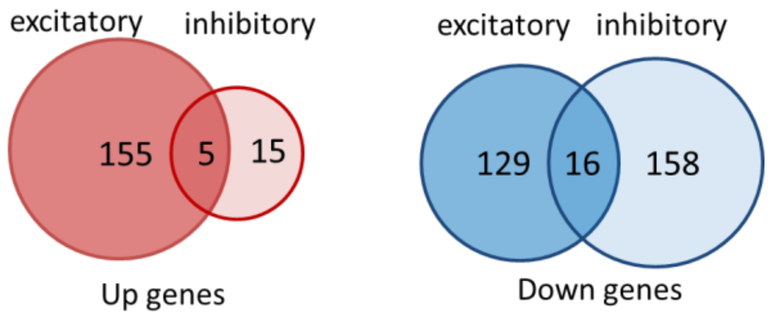


**Figure S7:** Venn diagram of consensus CI-DEGs in excitatory neuron and inhibitory neuron.

| **1 unit of tissue** | Absolute #Neuron | Absolute #Glia | Absolute mRNA | **Sent to sequencer** | Sequenced mRNA | Estimated #Neuron | Estimated #Glia |
|---|---|---|---|---|---|---|---|
| Control | 80 cells (160λ mRNA) | 20 cells (20λ mRNA) | 180λ | Control | 180λ | 80 cells | 20 cells |
| Case1: AD | 40 cells (80λ mRNA) | 20 cells (20λ mRNA) | 100λ | Case1: AD | 180λ | 72 cells | 36 cells |
| Case2: AD | 60 cells (120λ mRNA) | 30 cells (30λ mRNA) | 150λ | Case2: AD | 180λ | 72 cells | 36 cells |

**Figure S8:** Illustration of absolute change of cell proportion (blue panel) and relative change of cell proportion (pink panel) in a bulk RNAseq experiment. This is a hypothetical experiment to illustrate the effect of library preparation step on cell type proportions. For the ease of description, assume there are only two cell types – neurons and glia; assume the total mRNA amount of a glial cell is λ, and the total mRNA amount of a neuronal cell is 2λ on average. In the above scenario there is one control and two AD samples (Case 1 and Case 2). We assume that the cell counts in the Control are baseline (i.e. no changes due to disease). In Case 1, half of the neuronal cells were lost while the number of glia cells remains the same. In other words, the absolute proportion of neuronal cell loss is 50%. In a typical bulk RNAseq approach, the library preparation involves a normalization step (TruSeq® RNA Sample Preparation v2 Guide). The goal of this step is to send similar amounts of cDNAs to the sequencer. After this step, the (relative) neuronal loss detected in Case 1 is 10% that of the Control. In Case 2, one quarter of neuronal cells are lost and and glial cell increase by 50% compared to the Control (absolute changes). However, after the cDNA normalization step, the (relative) neuronal loss that can be detected is about 10% that of the Control. Therefore, even though a bioinformatics approach could accurately estimate the number of cells of different cell types that were sequenced, it would be difficult to trace back to the absolute proportion of cell changes.

### References:


1. McKenzie, A.T. *et al.* Brain Cell Type Specific Gene Expression and Co-expression Network Architectures. *Scientific Reports* **8**, 8868 (2018).
2. Zhong, Y., Wan, Y.-W., Pang, K., Chow, L.M. & Liu, Z. Digital sorting of complex tissues for cell type-specific gene expression profiles. *BMC Bioinformatics* **14**, 89 (2013).
